# Orthogonal Acoustic Control of Gene Expression Using a Synthetic T7RNAP Rapa-Inducible Dimerization System

**DOI:** 10.64898/2026.06.19.733485

**Authors:** Haoming Zhang, Avi Schroeder, Peng Xu

## Abstract

Precise spatiotemporal regulation of engineered microbes remains a critical bottleneck in synthetic biology. While ultrasound is extensively utilized for imaging and drug delivery, its translation into bacterial chassis is hindered by the lack of stringent biochemical triggers. Here, we present a rationally designed, ultrasound-responsive hybrid molecular switch based on a strict acoustic-biochemical "AND-gate". We engineered a highly sensitive split-T7 RNA polymerase system, which the dimerization and subsequent gene transcription can only be triggered in the presence of both ultrasound and a PEG-modified rapamycin. By systematically optimizing the acoustic parameters, we deployed this spatiotemporal switch to dynamically regulate a microbial consortium. With *lysisE* suicide protein as the output module, we achieved precise and programmable tuning of bacterial population in a co-culture system. This acoustic gating strategy may provide a robust and versatile toolkit for complex microbiome engineering and dynamic biomanufacturing.

Graphic Abstract

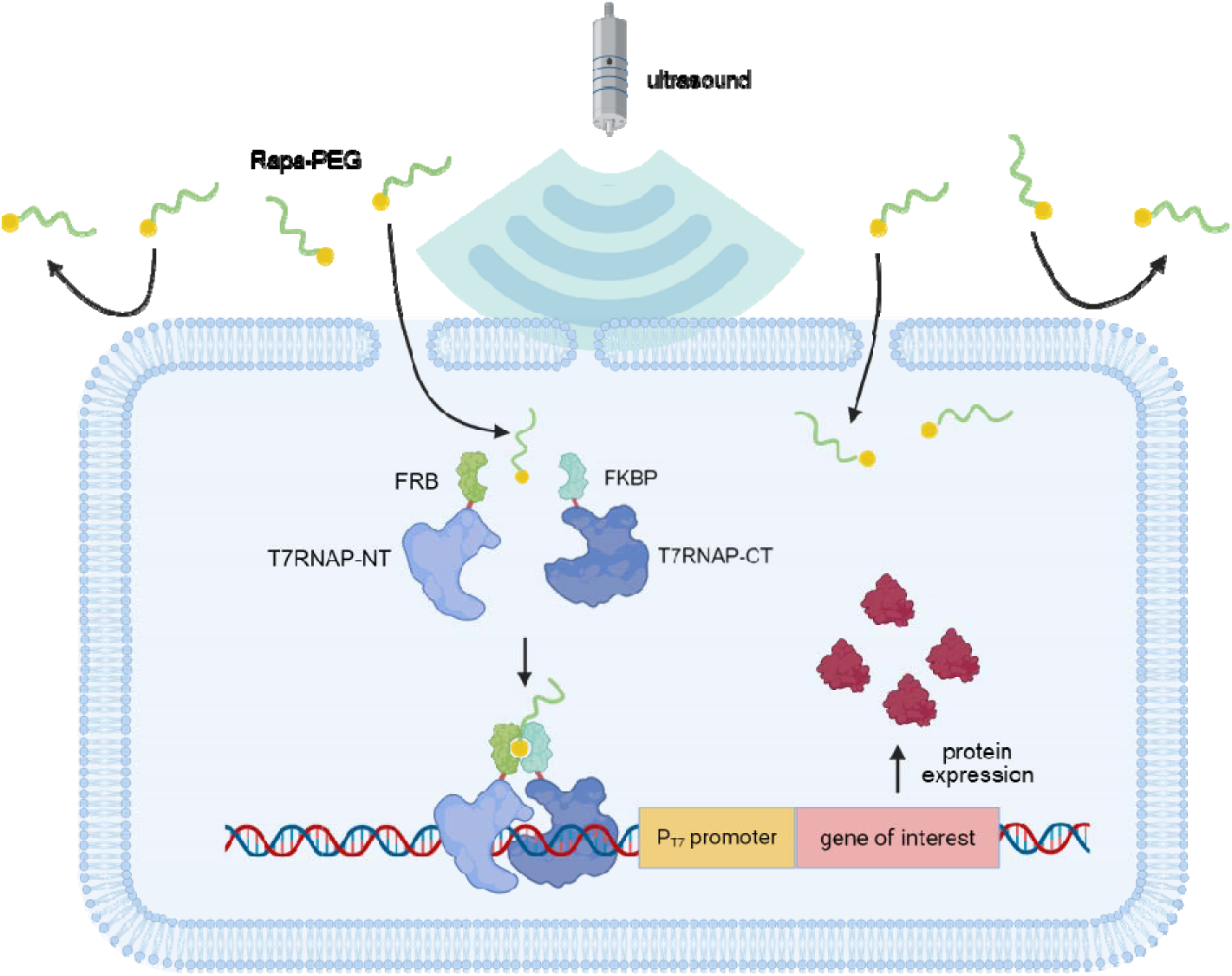

## Introduction

The dynamic regulation of microbial behavior lies at the heart of modern synthetic biology, enabling transformative applications ranging from adaptive biomanufacturing to smart live biotherapeutics^1,2^. As the field progresses from engineering isolated single strains to programming complex, multi-species microbial consortia, the ability to selectively and dynamically control specific sub-populations becomes paramount. In natural ecosystems, microbial consortia rely on intricate spatial and temporal resource allocation^3^. To mimic or engineer such complexity, synthetic biologists require orthogonal, highly tunable control switches that can be actuated with high spatiotemporal resolution within dense cultures or deep within host tissues.

Historically, the activation of synthetic genetic circuits has predominantly relied on small chemical regulators, such as isopropyl β-D-1-thiogalactopyranoside (IPTG), aTc, or arabinose. While effective for uniform bulk activation, chemical induction is fundamentally limited by slow diffusion dynamics, potential cellular toxicity, and significant off-target pleiotropy^4,5^. Moreover, once a chemical inducer is introduced into a closed system, gene expression cannot be easily halted or reversed, severely limiting its versatility in dynamic applications^6^. To overcome these limitations, recent advances in synthetic biology have introduced sophisticated control modalities, including engineered vector platforms, kinetic regulators, RNA-targeting systems, and computational tools^7–11^. Among physical stimuli, optogenetics provides unparalleled, sub-second spatiotemporal resolution^12,13^. However, it’s in vivo and industrial utility is critically hampered by the shallow penetration depth of light due to intense scattering and absorption in dense microbial fermentations or mammalian tissues. Other physical modalities, such as thermogenetics and magnetogenetics, offer improved penetration but are often constrained by slow activation kinetics, potential heat-induced cellular toxicity, or the requirement for cumbersome localized hardware.

Ultrasound, on the other hand, has recently emerged as a highly attractive modality for biological actuation. Widely utilized in diagnostic imaging and therapeutics^14–16^, ultrasound is non-invasive, cost-effective, and uniquely capable of penetrating deep into tissues (up to tens of centimeters) while maintaining millimeter-scale spatial focus^17^. Despite these profound advantages, the application of ultrasound for orthogonal gene expression control remains underexplored. Unlike optogenetic systems that rely on native photoreceptors, no naturally occurring transcriptional factors directly respond to acoustic waves. To bridge this gap, pioneering work in "sonogenetics" has explored using ultrasound to trigger stress-related promoters that respond to heat, protein aggregation, or membrane damage^18–20^. However, these systems rely on native *E. coli* pathways, making them highly susceptible to interference from the host’s endogenous regulatory networks and metabolic fluctuations. Furthermore, these endogenous stress promoters typically yield much lower expression levels compared to robust orthogonal promoters (e.g., T7) and inherently lack strict orthogonality. Alternative approaches have utilized acoustic radiation force to activate heterologous mechanosensitive channels (e.g., Piezo1 or MscL)^21,22^; however, these large, membrane-integrated receptors often suffer from high basal leakiness due to inherent membrane tension fluctuations and are challenging to reliably port into diverse bacterial chassis.

In this work, we propose a fundamentally distinct, physicochemical paradigm for acoustic control: size-dependent acoustic gating via sonoporation. Rather than engineering complex mechanosensitive proteins or hijacking non-orthogonal stress pathways, we focused on physically restricting the cellular entry of a highly potent, membrane-permeable chemical inducer (rapamycin) by conjugating it to a bulky, highly hydrophilic polyethylene glycol (PEG) chain. We hypothesized that this macromolecular modification (Rapa-PEG) would sterically and energetically prohibit the inducer from passively crossing the intact bacterial lipid bilayer under normal physiological conditions. Cellular entry—and subsequent robust gene activation—would exclusively occur during the transient, localized membrane fluidization and nanoscale pore formation (sonoporation) uniquely triggered by acoustic cavitation^23^.

To realize this concept, we engineered an ultrasensitive, low-noise genetic amplifier based on a split-T7 RNA polymerase biosensor, establishing a strict chemically induced dimerization (CID) dependency^24^. We synthesized a library of Rapa-PEG conjugates and validated their structural bio-orthogonality in a membrane-free cell-free (TX-TL) system, definitively decoupling their physical membrane impermeability from their intrinsic biological binding affinity. Through systematic in vivo screening, we identified an optimal "Goldilocks" conjugate (Rapa-PEG1000) that exhibits negligible basal leakage yet robust, ultrasound-dependent transcriptional activation, effectively functioning as a strict physical-chemical "AND-gate". Finally, we deployed this ultrasound-responsive switch to control a "suicide" module (*lysisE*) within a synthetic microbial consortium. By delivering targeted acoustic pulses, we achieved the selective ablation of an engineered strain, liberating metabolic resources and successfully driving a complete, programmable inversion of the population ratio. This robust, orthogonal, and modular strategy bridges the gap between acoustic physics and synthetic biology, providing a powerful new toolkit for non-invasive, spatiotemporally precise microbiome engineering.

## Results

### Engineering a High-Performance Split-T7 Biosensor

To establish an orthogonal and highly robust genetic amplifier for acoustic control, we first engineered a chemically inducible split-T7 RNA polymerase (T7RNAP) biosensor. T7RNAP was chosen for its exceptional transcriptional processivity and strict orthogonality to the endogenous regulatory networks of the host *E. coli*. We utilized a protein-fragmentation strategy, wherein the functional T7RNAP was divided into two inactive domains and subsequently genetically fused to the chemically induced dimerization (CID) pair: FK506-binding protein (FKBP) and the FKBP-rapamycin binding domain (FRB)^24,25^. In this rationally designed configuration, the presence of the small molecule rapamycin acts as a molecular bridge, inducing the heterodimerization of FKBP and FRB. This spatial proximity seamlessly reconstitutes the split-T7RNAP, thereby activating the transcription of the downstream mCherry reporter gene driven by the *P_T7_* promoter (**Figure 1A**)^26^.

**Figure 1.**
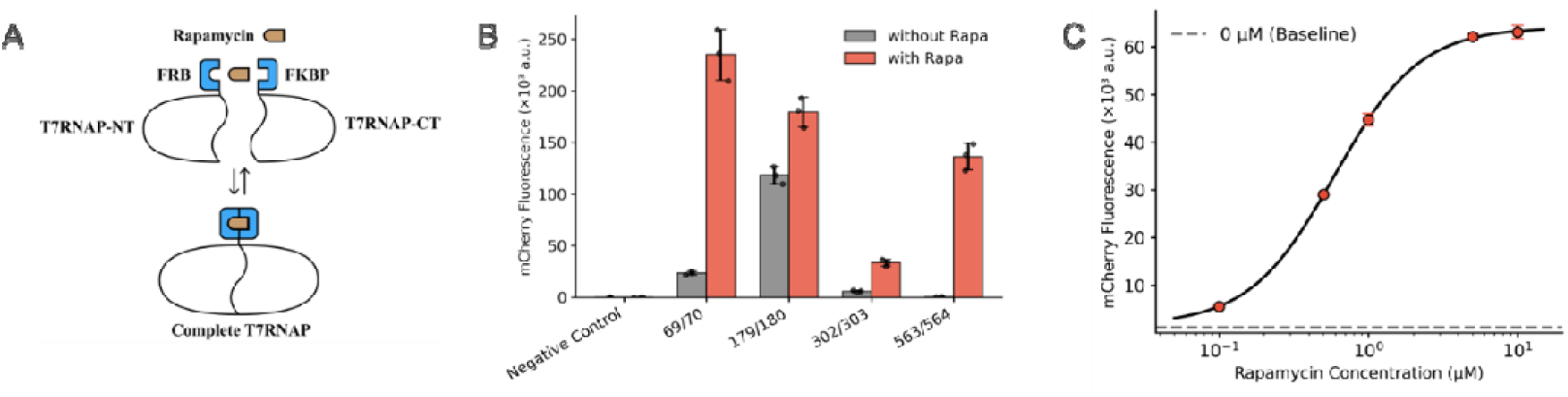
Design, screening, and characterization of the highly stringent split-T7 biosensor. (A) Schematic illustration of the chemically inducible split-T7 RNA polymerase (T7RNAP) system. The N- and C-terminal halves of T7RNAP are genetically fused to the FRB and FKBP domains, respectively, driving the expression of the downstream mCherry reporter via the P_T7_ promoter upon rapamycin addition. (B) Systematic screening of four different T7RNAP split variants (truncated at amino acids 69/70, 179/180, 302/303, and 563/564) in the presence (+Rapa) or absence (-Rapa) of the inducer. (C) Dose-response curve of the optimized 563/564 split-T7 biosensor under varying concentrations of native rapamycin.

Optimization of the split site is a critical engineering parameter to minimize spontaneous self-assembly (basal leakiness) while maximizing the transcriptional output upon induction^27^. Guided by previous optogenetic structural studies, we constructed and systematically screened a library of split-T7 variants truncated at highly permissive sites (amino acid positions 69/70, 179/180, 302/303, and 563/564). As illustrated in **Figure 1B**, the specific truncation site profoundly dictated the biosensor’s performance. The split at position 179/180 exhibited the highest absolute expression level; however, it suffered from severe basal leakiness (yielding substantial fluorescence even without rapamycin), rendering it unsuitable for precise spatial-temporal regulation. In sharp contrast, the split at position 563/564 demonstrated an exceptional signal-to-noise ratio. It maintained a remarkably low basal background in the absence of the inducer, while achieving a robust, nearly 280-fold activation upon the addition of rapamycin. Therefore, the 563/564 split configuration was identified as the optimal architectural scaffold and selected for all subsequent engineering. Furthermore, the effect of varying the fusion orientations and linker compositions on the biosensor performance was also evaluated, confirming the current architectural design as optimal (**Figure S1**).

To further define the operational dynamics of this optimized biosensor, we mapped its dose-response kinetics. We first monitored the time-course mCherry expression kinetics under varying concentrations of rapamycin, observing that the system consistently reached a steady-state transcription plateau approximately 8-10 hours post-induction (**Figure S2**). Utilizing this steady-state fluorescence data, we plotted the dose-response curve for the 563/564 split-T7 construct exposed to a rapamycin gradient ranging from 0.1 to 10 µM (**Figure 1C**). The system exhibited a highly sensitive, dose-dependent sigmoidal activation curve. The mCherry fluorescence signal increased sharply and approached saturation at concentrations above 5 µM. This saturation behavior at elevated concentrations is likely attributed to the intrinsic poor aqueous solubility of rapamycin, which limits its effective intracellular availability and promotes extracellular aggregation. Nevertheless, this highly sensitive and tightly regulated split-T7 system establishes a reliable, low-noise biological chassis, laying the fundamental groundwork for the subsequent integration of our ultrasound-responsive molecular switch.

### Rational Design, Synthesis, and Cell-Free Validation of the Molecular Switch

As established in the previous section, native rapamycin is a highly potent inducer; however, its inherent lipophilicity allows it to freely and passively diffuse across the bacterial cell membrane. This uncontrolled entry renders native rapamycin fundamentally unsuitable for spatiotemporally gated activation. To engineer an ultrasound-responsive trigger, our core design rationale was to structurally impede this passive diffusion, thereby restricting cellular entry exclusively to periods of acoustic membrane disruption. Ultrasound has been shown to temporarily increase cell permeability by creating transient nanoscale pores in the cell membrane, a process known as sonoporation^23,28^. This effect is primarily driven by the mechanical forces generated by acoustic waves, which cause microbubbles in the surrounding medium to oscillate and collapse, leading to localized increases in pressure and lipid bilayer mobility^29,30^. These temporary pores allow the controlled entry of target compounds without significantly affecting overall cell viability.

The *E. coli* cell membrane is composed of a lipid bilayer that is inherently hydrophobic, severely restricting the passive transit of large hydrophilic molecules^31^. Based on this property, we hypothesized that covalently fusing rapamycin to a bulky, highly hydrophilic polymer could effectively minimize its passive diffusion. Polyethylene glycol (PEG) was selected as the optimal modifier due to its extensive use in pharmacology, extreme hydrophilicity, and easily tunable molecular weight.

To identify an appropriate modification site without destroying the molecule’s chemically induced dimerization (CID) function, we examined the X-ray crystal structure of the FKBP-rapamycin-FRB ternary complex (PDB ID: 1FAP)^32^ (**Figure 2A**). Structural analysis reveals that the C40-hydroxyl group on the rapamycin cyclohexyl ring is highly solvent-exposed and prominently oriented outward, strictly away from the critical protein-ligand interaction interfaces.

**Figure 2.**
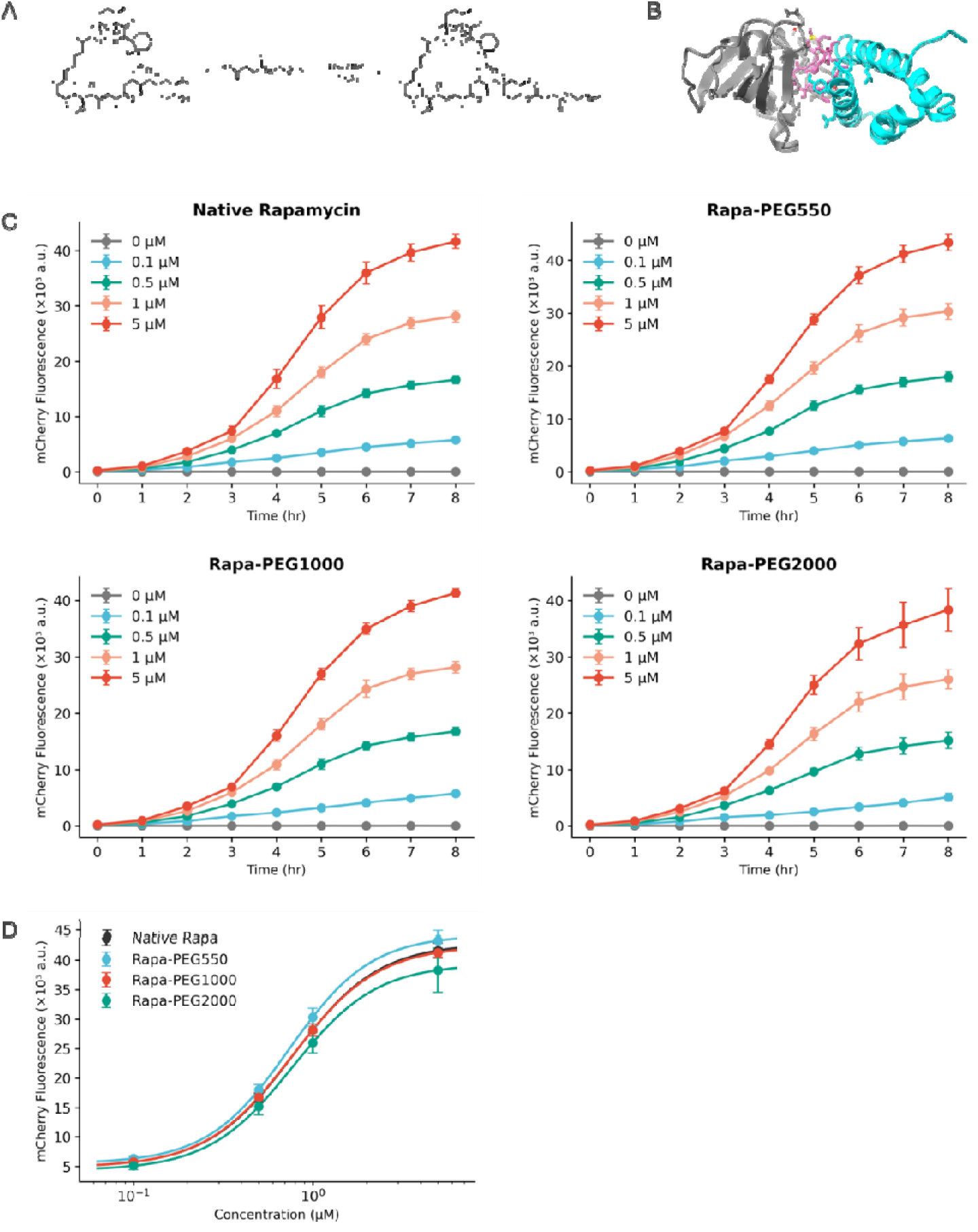
Rational design, chemical synthesis, and cell-free validation of the size-excluded Rapa-PEG conjugates. (A) Schematic of the chemical synthesis route. Methoxy-polyethylene glycol-carboxylic acids (mPEG-COOH) of varying molecular weights (Mw = 550, 1000, and 2000 Da) were conjugated to the C40-OH position of native rapamycin via an EDC/DMAP-mediated esterification reaction to yield the Rapa-PEG variants. (B) Structural analysis based on the X-ray crystal structure of the FKBP12-rapamycin-FRB ternary complex (PDB ID: 1FAP). The 3D model highlights that the C40-hydroxyl group of rapamycin (highlighted in yellow) is highly solvent-exposed and oriented away from the critical protein-binding interfaces. (C) Cell-free transcription-translation (TX-TL) assay comparing mCherry expression levels induced by native rapamycin and the synthesized Rapa-PEG variants in a membrane-free environment. (D) Normalized dose-response curves for native rapamycin and the varying Rapa-PEG conjugates in the cell-free system.

Based on this structural geometry, we hypothesized that the site-specific conjugation of bulky, hydrophilic polymers at the C40 position would freely extend into the surrounding aqueous environment without incurring steric clashes that disrupt FKBP-FRB heterodimerization. Simultaneously, this modification would effectively transform the membrane-permeable small molecule into a massive, heavily hydrated macromolecule, energetically and sterically prohibiting it from crossing the intact lipid bilayer.

To execute this design, we covalently conjugated native rapamycin to biologically inert polyethylene glycol (mPEG-COOH) chains of varying molecular weights (Mw = 550, 1000, and 2000 Da) via an esterification reaction directed at the C40-OH position (**Figure 2B**) according to a previously reported method^33^. PEG is highly hydrophilic and strongly coordinates with surrounding water molecules, which dramatically expands the hydrodynamic radius of the conjugate. The successful synthesis and high purity of the resulting Rapa-PEG variants were rigorously confirmed via ^1^H NMR and TLC (**Figure S3**).

A critical prerequisite for this acoustic gating strategy was to confirm our hypothesis that PEGylation does not compromise the innate binding affinity of the rapamycin core. To completely decouple the intrinsic biological activity of the synthesized conjugates from their altered cellular permeability, we systematically evaluated their performance in a membrane-free, cell-free transcription-translation (TX-TL) system harboring the split-T7 biosensor. The time-course kinetic profiles demonstrated that all Rapa-PEG variants rapidly initiated mCherry expression, reaching steady-state plateaus within 8 hours (**Figures 2C** and **2D**).

Strikingly, steady-state dose-response analysis revealed that the conjugation of PEG chains—regardless of their length or hydrodynamic volume—did not hinder transcriptional output. All three variants induced robust reporter expression at levels highly comparable to unmodified native rapamycin, exhibiting nearly identical sigmoidal activation profiles with similar EC_50_ values (**Figure 2E**).

This robust, size-independent induction *in vitro* provides definitive, mechanistically grounded proof of the retained structural bio-orthogonality. It unambiguously establishes that any functional failure of larger conjugates (e.g., Rapa-PEG2000) observed in living cells would be strictly attributable to physical, size-dependent membrane exclusion, rather than a loss of chemical binding affinity. This perfect cell-free baseline thereby lays a rigorous, irrefutable foundation for exploring ultrasound-mediated membrane permeabilization in subsequent in vivo studies.

### Size-Dependent Acoustic Gating and Spatiotemporal Modulation

Having established the split-T7 biosensor and confirmed the bio-orthogonality of PEGylated rapamycin *in vitro*, we next sought to achieve ultrasound-dependent gene activation *in vivo*. The underlying principle relies on transient acoustic cavitation (sonoporation) in the bulk liquid medium, which temporarily increases the permeability of the bacterial cell envelope, allowing otherwise impermeable macromolecules to enter the cytoplasm^18^.

To execute this, bacteria were grown in shake flasks to the logarithmic phase (OD_600_ 0.8–1.0) before being subjected to ultrasound in the presence of 5 µM Rapa-PEG. To minimize the negative impacts of physical stress, such as overheating, the acoustic treatment was applied in cycles consisting of one minute of pulsed ultrasound followed by three minutes of recovery in a shaker. This 3-minute shaking interval is mechanistically crucial for three reasons: firstly, it ensures the recovery of cell viability from the ultrasonic challenge; secondly, the vigorous mixing homogenizes the Rapa-PEG molecules, maximizing their probability of encountering the permeabilized regions of the cell membrane; and finally, it rapidly regenerates dissolved oxygen and the gas microbubbles required for the next round of cavitation-induced membrane disruption.

We hypothesized that the gating efficiency—defined by low basal leakage and high ultrasound-induced activation—would be strictly dependent on the hydrodynamic radius of the Rapa-PEG conjugates. To test this size-dependent gating mechanism, we evaluated the *in vivo* performance of Rapa-PEG variants with different PEG chain lengths (Mw 550, 1000, and 2000 Da). The results revealed a striking size-activity relationship (**Figure 3A**). The smallest conjugate, Rapa-PEG550, exhibited unacceptably high basal leakiness even in the absence of ultrasound, suggesting that its relatively small hydrodynamic volume allowed it to passively diffuse across the bacterial membrane. Conversely, the largest conjugate, Rapa-PEG2000, showed minimal basal leakage but failed to induce substantial gene expression even upon ultrasound stimulation, indicating that its massive steric bulk hindered cellular entry despite sonoporation-induced membrane fluidization. Remarkably, Rapa-PEG1000 hit the optimal "Goldilocks" zone. It maintained a strict low basal expression level in the unperturbed state, while triggering a robust activation of mCherry exclusively upon ultrasound application. Therefore, Rapa-PEG1000 was selected as the designated molecular switch for all subsequent experiments.

**Figure 3.**
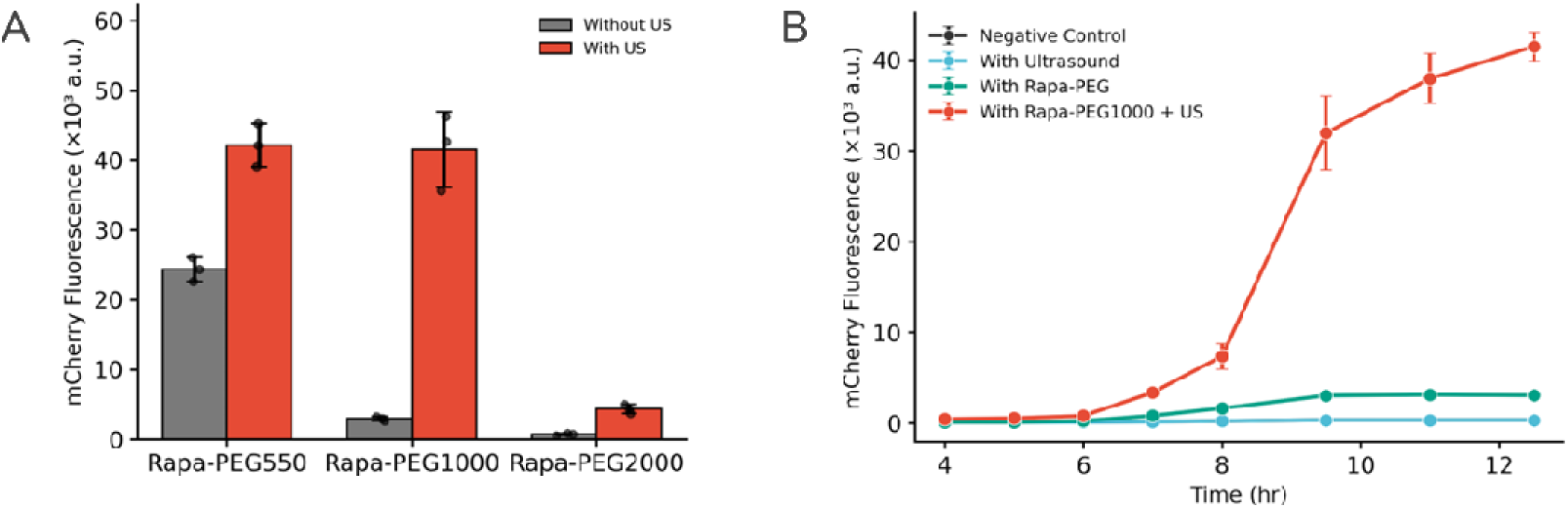
In vivo validation of the size-dependent acoustic gating mechanism. (A) In vivo screening of the Rapa-PEG conjugates of varying molecular weights (Mw = 550, 1000, and 2000 Da) in engineered E. coli in the presence (+US) or absence (-US) of ultrasound treatment. (B) Real-time in vivo fluorescence kinetics of the engineered bacteria under different conditions: untreated (negative control, black line), ultrasound alone (blue line), Rapa-PEG1000 alone (green line), and the combination of Rapa-PEG1000 and ultrasound (red line).

A key advantage of integrating physical stimuli like ultrasound into synthetic biology is the capability to achieve non-invasive, highly tunable spatiotemporal control. Using the optimized Rapa-PEG1000 switch (maintained at a constant concentration of 5 µM), we extensively characterized the system’s dynamic range. First, we monitored the time-course kinetics of the acoustic induction. As shown in **Figure 3B**, the control groups—including cells exposed to ultrasound alone and cells incubated with Rapa-PEG1000 without ultrasound—showed negligible mCherry expression over the 11-hour period. In stark contrast, cells treated with both Rapa-PEG1000 and ultrasound exhibited a rapid and sustained increase in fluorescence, reaching a steady state after 8 hours. This strict "AND-gate" logic definitively proves that acoustic stimulation is the indispensable trigger for the engineered molecular switch.

To further demonstrate the programmability of this hybrid system, we systematically mapped the gene expression output against varying acoustic parameters. By fixing the sonication duration and varying the ultrasound frequency (0, 5, 10, and 20 kHz), we observed a distinct threshold for effective sonoporation (**Figure 4A**). Compared to the untreated control, a 5 kHz treatment resulted in only a marginal increase in mCherry signal, indicating that the acoustic energy was insufficient to effectively agitate the membrane. However, when the frequency was elevated to 10 kHz, the permeability increased dramatically, leading to a substantial surge in fluorescence. A frequency of 20 kHz further maximized the transcriptional output. We intentionally capped the maximum frequency at 20 kHz based on previous biophysical studies demonstrating that *E. coli* can withstand this specific frequency without suffering severe, irreversible structural damage or catastrophic loss of viability^34^.

**Figure 4.**
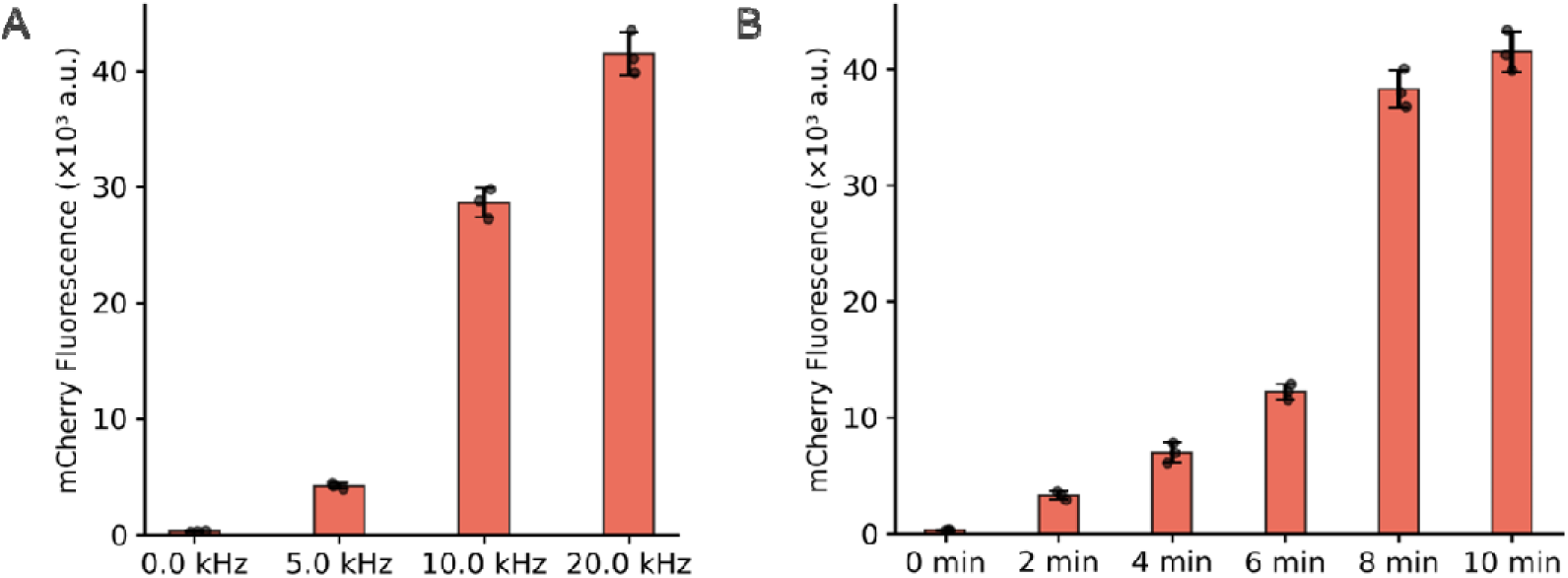
Systematic optimization of acoustic actuation parameters. (A) Evaluation of ultrasound frequency on biosensor activation. Engineered bacterial cultures supplemented with Rapa-PEG1000 were subjected to acoustic treatments at various frequencies (5, 10, and 20 kHz) using a fixed exposure duration of 10 minutes and compared to an untreated control (0 kHz). (B) Evaluation of ultrasound exposure duration. Using a fixed frequency of 20 kHz, the duration of the pulsed ultrasound exposure was systematically varied from 0 to 10 minutes to monitor the corresponding intracellular accumulation of mCherry.

Subsequently, holding the frequency constant at 20 kHz, we systematically varied the total ultrasound duration from 0 to 10 minutes (**Figure 4B**). In general, the mCherry signal increased alongside the treatment time, but this growth was notably non-linear. The fluorescence output at 8 minutes was drastically higher than at 6 minutes, reflecting a compounding effect as progressively more cells in the actively growing culture were effectively permeabilized. However, extending the treatment to 10 minutes yielded only a marginal enhancement over the 8-minute mark. This plateau suggests that the sonoporation efficiency had approached its theoretical upper limit, or that prolonged physical stress began to counterbalance the expression gains.

Collectively, these results demonstrate that by rationally adjusting ultrasound parameters, our system allows for the precise, user-defined titration of gene expression, effectively circumventing the slow dynamics and irreversibility often associated with traditional chemical inducers. This robust spatiotemporal tunability provides a versatile tool for complex synthetic biology applications, such as dynamic microbial consortium control.

### Programmable Population Control in Synthetic Consortia

Having demonstrated the precise spatiotemporal tunability of our ultrasound-responsive switch, we next sought to apply this system to tackle a major challenge in synthetic biology: the dynamic regulation of sub-populations within a microbial consortium^35,36^. Typically, the initial strain ratio is fixed at the time of inoculation, but it frequently deviates over time as strains compete for limited nutrients, undermining the intended balance and limiting the feasibility of large-scale biomanufacturing^37,38^. We hypothesized that our acoustic-gating system could dynamically intervene and regulate strain ratios *in situ*.

To engineer a "suicide" module, we placed the *lysisE* gene under the control of the *P_T7_* promoter (**Figure S4**). LysisE is a phage-derived protein whose accumulation abruptly disrupts fatty acid synthesis, leading to catastrophic cell envelope failure and lysis^39^. Initially, plasmid construction proved exceedingly difficult due to the acute toxicity of LysisE; practically no colonies could be recovered post-transformation. This hurdle was resolved by appending a C-terminal LVA degradation tag to the LysisE protein. This modification substantially decreased its cellular residence time, preventing lethal accumulation caused by inevitable minor basal leakiness from the *P_T7_* promoter.

Before deploying the co-culture system, we first validated the acoustic-triggered selective lysis efficiency in a monoculture. As depicted in **Figures 5B**, the engineered strain exhibited normal growth kinetics in the uninduced state. However, upon the synergistic application of ultrasound and the Rapa-PEG1000 switch, the optical density (OD_600_) of the culture plummeted rapidly after a brief lag phase. Crucially, to rule out the possibility of non-specific mechanical damage caused by acoustic cavitation, a wild-type (WT) *E. coli* control was subjected to the maximum ultrasound duration (8 min). The WT strain exhibited an unperturbed growth trajectory virtually identical to the uninduced control, definitively proving that the observed population crash is entirely driven by our engineered genetic suicide switch rather than generic physical stress.

**Figure 5.**
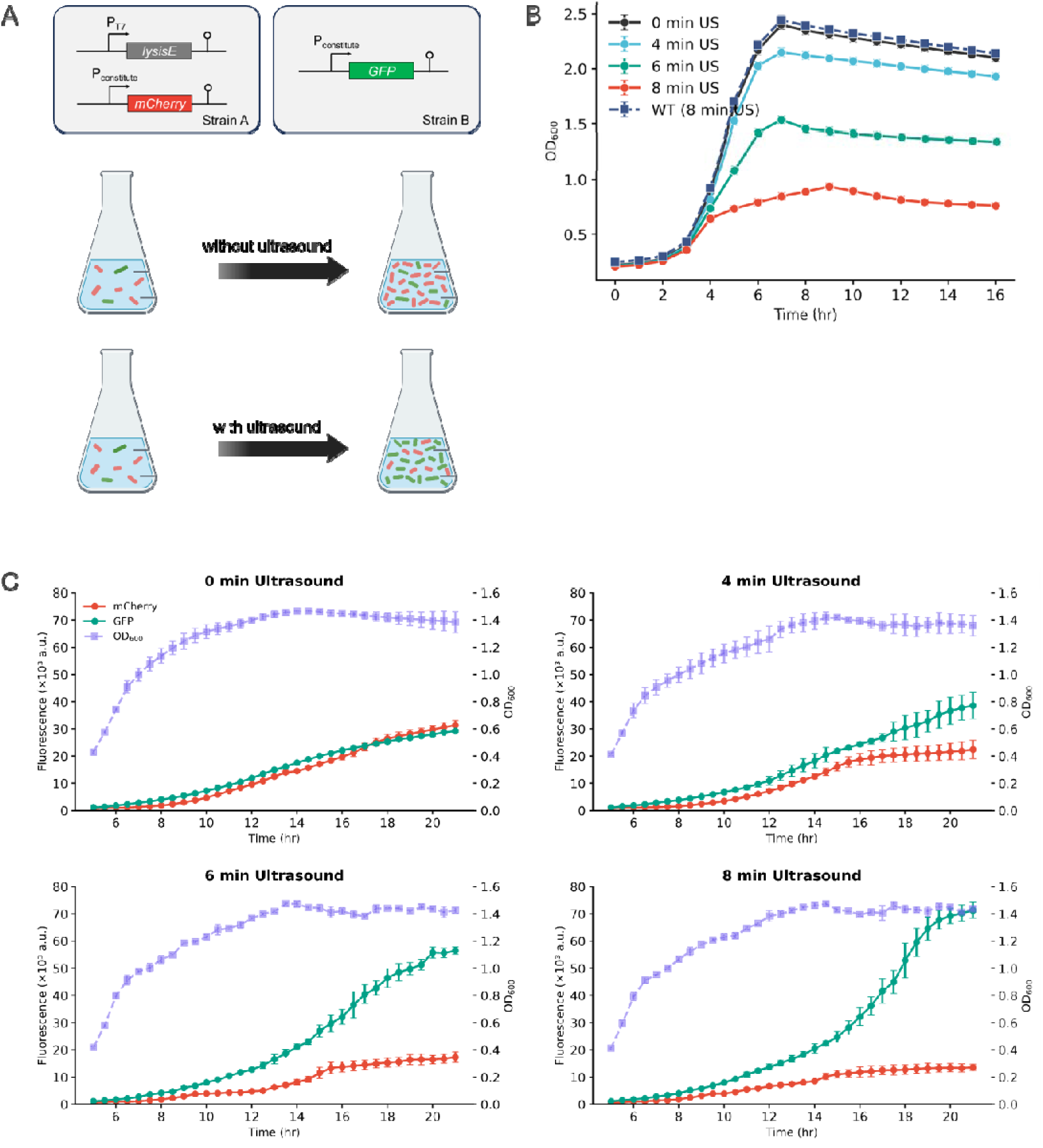
Programmable population control and targeted species ablation in a synthetic microbial consortium. (A) Schematic illustration of the synthetic consortium strategy, consisting of an acoustic-responsive lysis strain (red) and an orthogonal, constitutively active survivor strain (green). (B) Validation of ultrasound-triggered cell lysis in monoculture. Optical density (OD_600_) profiles of the engineered "suicide" strain (harboring the acoustic-gated *lysisE* module) subjected to varying durations of ultrasound treatment (0, 4, 6, and 8 min) in the presence of Rapa-PEG1000. A wild-type (*WT*) E. coli control subjected to the maximum 8 min ultrasound treatment is included for comparison. (C) Dynamic tracking of population inversion in co-culture. The two strains were co-cultured at an initial 3:1 ratio (mCherry:GFP) and subjected to varying ultrasound durations (0, 4, 6, and 8 min at 20 kHz). Temporal profiles of absolute fluorescence (mCherry, red solid line; GFP, green solid line) and total cell density (OD_600_, purple dashed line) were monitored over a 20-hour period.

We then constructed a competitive synthetic consortium comprising two distinct populations (**Figure 5A**): the acoustic-responsive lysis strain (labeled with mCherry) and an acoustically orthogonal, constitutively active survivor strain (labeled with GFP). The co-culture was inoculated at an initial ratio of 3:1 (mCherry:GFP). It is worth noting that without ultrasound, the baseline macroscopic fluorescence outputs of mCherry and GFP appeared visually comparable at early stages (**Figure 5C, 0 min group**). This is because equivalent protein expression levels do not necessarily yield identical fluorescence intensities due to inherent differences in quantum yields, excitation/emission efficiencies, and equipment sensitivity profiles. In a nutrient-limited batch culture without acoustic intervention, the population remained relatively stable and coexisted throughout the cultivation period.

Strikingly, the application of ultrasound fundamentally reprogrammed the consortium’s trajectory. As the ultrasound duration increased, the growth of the mCherry population was forcefully arrested and subsequently depleted. This targeted ablation liberated critical nutrients and spatial niches, which were rapidly scavenged by the GFP-expressing survivor strain. Consequently, the total OD_600_ of the consortium remained relatively constant, but the GFP fluorescence intensity surged, achieving a near-complete population inversion by the 10-hour mark.

To visually confirm this selective lysis mechanism and the ensuing population shift in situ, we performed time-lapse confocal laser scanning microscopy (CLSM) across varying ultrasound exposure durations (0, 4, 6, and 8 minutes at 20 kHz) (**Figure 6A**). At the initial 0-hour time point, all groups displayed a similar dual-color coexistence profile corresponding to the inoculation ratio. By the 4-hour mark, distinct morphological and population differences began to emerge in a strictly ultrasound-dose-dependent manner as *lysisE* expression initiated. After 8 hours, the divergence was absolute: the control group without ultrasound (0 min) maintained stable coexistence, whereas the groups subjected to longer acoustic treatments exhibited a progressive, dose-dependent eradication of the mCherry population. Specifically, the 8-minute actuated group transitioned into an overwhelmingly green-dominated community, visually corroborating the quantitative macroscopic data.

**Figure 6.**
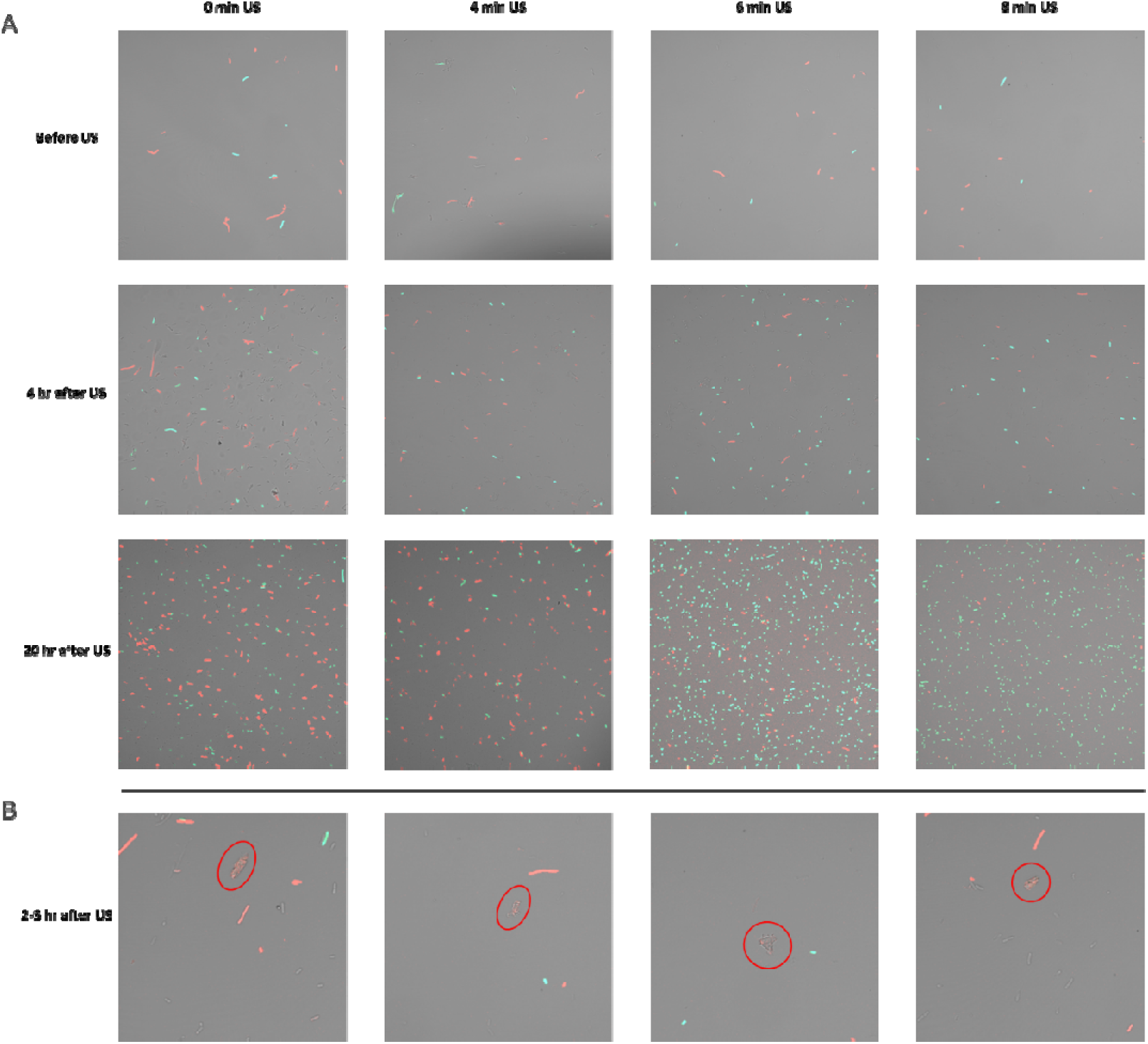
Single-cell and macroscopic tracking of acoustic-programmed population inversion in a synthetic microbial consortium. (A) Time-course confocal laser scanning microscopy (CLSM) imaging of the synthetic consortium. The co-culture was subjected to varying durations of ultrasound treatment (0, 4, 6, and 8 min at 20 kHz) in the presence of Rapa-PEG1000 and monitored at 0, 4, and 8 hours post-actuation. (B) High-resolution CLSM observation capturing individual mCherry-expressing *E. coli* cells in the active process of lysisE-mediated membrane disruption, evidenced by localized swelling and extracellular diffusion of the red fluorescent reporter.

More importantly, high-magnification imaging at an early intermediate stage (2–3 hours post-sonication) captured the transient biophysical events governing this shift. These images revealed distinct morphological hallmarks of programmed cell death exclusively in the mCherry-labeled strain (**Figure 6B**). Localized swelling of the cell envelope and the characteristic extracellular diffusion of red fluorescence provided direct, visual evidence of lysisE-mediated membrane disruption and subsequent protein leakage. Together, these multiscale validations—from bulk kinetic assays to single-cell morphological observations—firmly establish our ultrasound-gated molecular switch as a robust and non-invasive tool for dynamic microbiome engineering.

## Discussion

The precise spatiotemporal regulation of engineered cellular functions remains one of the most formidable challenges in synthetic biology. While chemical induction is ubiquitous, its reliance on passive diffusion fundamentally precludes localized control and is plagued by severe off-target pleiotropy. Conversely, physical triggers like light (optogenetics) offer superior temporal resolution but suffer from catastrophic signal attenuation in deep tissues or dense industrial cultures. In this study, we addressed these critical bottlenecks by conceptualizing and validating a rationally designed, ultrasound-gated molecular switch. By transforming a highly potent but freely diffusing chemical inducer into a sterically excluded "physical-chemical AND-gate," our system harnesses the deep-tissue penetrability of acoustic waves while strictly preserving the absolute orthogonality and modularity of genetic biosensors.

The core elegance of our design lies in the structural decoupling of biological binding affinity from physical membrane permeability. Traditional approaches to sonogenetics often rely on the overexpression of mechanosensitive channels (e.g., MscL or Piezo1) to perceive acoustic waves^22,40^. However, these systems frequently suffer from high basal leakiness due to inherent membrane tension fluctuations and lack true molecular orthogonality in bacterial chassis^21,41^. Instead of altering the cellular hardware, we engineered the inducer itself. Guided by the crystal structure of the FKBP-rapamycin-FRB complex, our site-specific C40-PEGylation generated a "Goldilocks" probe (Rapa-PEG1000)—massive enough to be completely excluded by the intact E. coli lipid bilayer, yet fully capable of bridging the CID components once introduced into the cytoplasm via transient sonoporation. The perfect alignment of our cell-free (TX-TL) validations with the in vivo acoustic actuation data compellingly demonstrates that synthetic structural biology can be utilized to rationally program physical transport constraints without compromising macromolecular recognition.

Beyond the fundamental proof-of-concept, we demonstrated the profound utility of this acoustic switch in resolving a major limitation in microbial consortium engineering. In nature, microbial communities thrive on complex spatial and temporal resource allocations; however, synthetic co-cultures in batch fermentation often succumb to competitive exclusion, where minor differences in growth rates inevitably lead to the unintended collapse of the slower-growing strain^42^. By genetically hardwiring our ultrasound switch to the lysisE suicide module, we achieved the targeted ablation of a specific sub-population, driving a complete and programmable population inversion.

This non-invasive intervention capability opens highly promising avenues for advanced biomanufacturing and metabolic engineering. In complex biosynthetic pathways involving metabolic division of labor, co-cultures are often employed where an upstream strain synthesizes an intermediate precursor, which is subsequently converted into the final product by a downstream strain. Determining and maintaining the optimal population ratio to prevent precursor bottlenecking or metabolic burden is notoriously difficult. Our acoustic gating system could be seamlessly deployed in such bioreactors: once the upstream strain has accumulated sufficient precursors, a precisely tuned ultrasound pulse could trigger its programmed lysis. This would simultaneously halt unnecessary nutrient consumption, eliminate competitive metabolic burden, and release the intracellular precursors into the bulk medium, making them entirely accessible for the downstream biotransformation by the survivor strain.

While our approach shows immense promise, we acknowledge certain physical parameters that warrant future optimization. Transient acoustic cavitation fundamentally introduces mechanical stress and localized micro-streaming, which could perturb the global transcriptomic profile of the host cells if the acoustic dosage is excessively high^43^. Although we mitigated non-specific toxicity by adopting a pulsed ultrasound regimen (e.g., 1 min actuation followed by 3 min recovery) and limiting the frequency to an empirically safe 20 kHz threshold, the scaling of this technology to massive industrial fermenters will require rigorous acoustic mapping to ensure homogeneous cavitation fields. Furthermore, for future in vivo therapeutic applications (e.g., dynamic control of engineered probiotics in the mammalian gut), the integration of focused ultrasound (FUS) technologies will be crucial to achieve millimeter-level spatial resolution without causing collateral tissue hyperthermia^44^.

In conclusion, we have developed a highly versatile and programmable ultrasound-responsive genetic system in *E. coli*. By bridging physical sonoporation with synthetic molecular logic, we provide a robust paradigm for dynamic microbiome engineering. This non-invasive, deep-penetrating control strategy holds significant potential to advance both scalable metabolic biomanufacturing and the next generation of smart live biotherapeutics.

## Materials and Methods

### Materials, Bacterial Strains, and Growth Conditions

*Escherichia coli* strain *DH5*α was utilized for all routine molecular cloning and plasmid construction. For functional characterization, monoculture assays, and co-culture experiments, *E. coli DH5*α (or a specified expression strain) was used as the host chassis to avoid interference from endogenous T7 RNA polymerase. Cells were routinely cultured in Luria-Bertani (LB) broth (10 g/L tryptone, 5 g/L yeast extract, 10 g/L NaCl) at 37 °C with continuous orbital shaking at 220 rpm. Solid media were prepared by supplementing LB broth with 1.5% (w/v) agar. Media were supplemented with ampicillin 100 µg/mL to maintain plasmids. Native rapamycin and polyethylene glycol (PEG) of varying molecular weights (Mw 550, 1000, and 2000 Da) were purchased from Sigma-Aldrich. Stock solutions of rapamycin and Rapa-PEG conjugates were prepared in dimethyl sulfoxide (DMSO). During induction, the final concentration of DMSO in the bacterial cultures was strictly maintained below 0.1% (v/v) to preclude any solvent-induced cytotoxicity.

### Plasmid Construction and Genetic Engineering

All recombinant plasmids were constructed utilizing either Gibson Assembly (NEBuilder HiFi DNA Assembly Master Mix, New England Biolabs) or standard restriction-ligation cloning. DNA sequences encoding the chemically induced dimerization (CID) domains (FKBP and FRB) and the phage-derived lysis gene (*lysisE*) were computationally codon-optimized for *E. coli* expression and synthesized *de novo* by Jiutian Gene Tech (Nanjing, China). The split-T7 RNA polymerase fragments (truncated at highly permissive sites, including 69/70, 179/180, 302/303, and 563/564) were amplified by polymerase chain reaction (PCR) utilizing Phusion Flash High-Fidelity DNA Polymerase (Thermo Fisher Scientific). Assembled plasmids were transformed into chemically competent *E. coli DH5*α. Transformants were selected on corresponding antibiotic plates and verified via colony PCR and Sanger sequencing (Genewiz, China) to ensure sequence integrity prior to downstream applications. Comprehensive plasmid maps and sequence details are provided in the Supporting Information.

### Synthesis of Rapa-PEG Conjugates

The Rapa-PEG conjugates (Rapa-PEG550, Rapa-PEG1000, and Rapa-PEG2000) were synthesized via an EDC/DMAP-mediated esterification reaction between the C40-hydroxyl group of rapamycin (RAPA) and the terminal carboxyl group of methoxy-PEG-carboxylic acid (mPEG-COOH). The final purified products were structurally validated by ^1^H NMR spectroscopy. Detailed synthetic procedures, purification steps, and structural characterization data are described in the Supplementary Methods.

### Cell-Free Transcription-Translation (TX-TL) Assay

To validate the structural bio-orthogonality of the synthesized conjugates without the confounding variable of the cell membrane, a commercial E. coli-based cell-free protein synthesis system (myTXTL, Arbor Biosciences) was employed. Plasmids encoding the split-T7 biosensor and the P_T7_-mCherry reporter were added to the TX-TL master mix at optimized concentrations. Rapa-PEG variants or native rapamycin were then added to reach final concentrations ranging from 0.1 to 5 µM. The reactions (10 µL volumes) were incubated at 30 °C in a 384-well black microplate. Fluorescence kinetics were monitored continuously using a multi-mode microplate reader.

### Acoustic Stimulation Apparatus and Parameters

For in vivo monoculture assays, bacterial cells at the exponential growth phase (OD600 ∼ 0.4–0.6) were supplemented with Rapa-PEG. Upon reaching an OD600 of 0.8–1.0, cultures were subjected to targeted ultrasound actuation using a customized 20 kHz probe-type sonicator operating under a pulsed duty cycle to prevent thermal damage. Optical density and mCherry fluorescence were recorded subsequently at 37 °C. Detailed physical parameters and the acoustic actuation setup are provided in the Supplementary Methods.

### *In Vivo* Gene Expression and Monoculture Lysis Assays

Bacterial overnight cultures were diluted 1:100 into fresh LB medium and incubated until reaching the exponential growth phase (OD_600_ ∼ 0.4–0.6). At this juncture, the optimized Rapa-PEG1000 inducer was supplemented to the cultures. The cells were allowed to further proliferate until reaching an OD_600_ of 0.8–1.0, at which point the predefined ultrasound treatment was administered. Following acoustic stimulation, 200 µL aliquots of the cultures were transferred into 96-well black, clear-bottom microplates (Corning). The microplates were incubated in a multimode microplate reader (e.g., Tecan Spark or BioTek Synergy) at 37 °C with continuous double-orbital shaking. Optical density (OD_600_) and fluorescence intensities were recorded automatically at specified intervals. mCherry fluorescence was acquired using an excitation of 580 nm and emission of 625 nm.

### Synthetic Consortium Co-Culture and Population Dynamics

For microbial consortium experiments, two distinct *E. coli* strains were engineered: a "lysis" strain constitutively expressing mCherry and harboring the acoustic-gated *lysisE* module, and a "survivor" strain constitutively expressing turboGFP. The two strains were inoculated into a single flask of nutrient-limited medium at an initial population ratio of 3:1 (mCherry:turboGFP). Rapa-PEG1000 was added, and the co-culture was subjected to targeted ultrasound treatment. Population dynamics were tracked over 10 hours by simultaneously measuring mCherry (Ex 580 nm / Em 625 nm) and turboGFP (Ex 482 nm / Em 502 nm) bulk fluorescence in a microplate reader. The fluorescence signals were normalized to the total OD_600_ to accurately reflect the relative abundance of each sub-population.

### Confocal Laser Scanning Microscopy (CLSM)

To obtain morphological evidence of acoustic-triggered cell lysis and to visualize the in situ spatiotemporal population shift, treated co-culture samples were analyzed via CLSM. At designated time points (e.g., 0 h, 2–3 h, 4 h, and 10 h post-ultrasound), 5 µL of the bacterial suspension was immobilized onto freshly prepared 1% (w/v) agarose pads or poly-L-lysine-coated glass-bottom dishes. Imaging was performed using a confocal microscope (Zeiss LSM 900) equipped with a 63× oil-immersion objective. turboGFP was excited with a 488 nm laser, and mCherry was excited with a 561 nm laser. To ensure unbiased comparative analysis, identical laser power, detector gain, and exposure settings were strictly maintained across all samples and time points. Post-acquisition image processing, including scale bar insertion and minor background adjustments, was performed utilizing ImageJ/Fiji software.

## Notes

### Competing Interest Statement

The authors have declared no competing interest.

