## Supplementary Information for "Orthogonal Acoustic Control of Gene Expression Using a Synthetic T7RNAP Rapa-Inducible Dimerization System"

**Supplementary Methods**

1. **Detailed Synthesis and Characterization of Rapa-PEG Conjugates**

An optimized 1:3 molar ratio of RAPA to mPEG-COOH was utilized for all syntheses to ensure high conjugation efficiency. Briefly, RAPA (0.3 mmol), the respective mPEG-COOH (Mw 550, 1000, or 2000 Da, 0.9 mmol), 1-(3-dimethylaminopropyl)-3-ethylcarbodiimide hydrochloride (EDC·HCl, 0.9 mmol), and 4-dimethylaminopyridine (DMAP, 0.09 mmol) were dissolved in 10 mL of anhydrous dichloromethane (DCM). The reaction mixture was stirred continuously for 3 hours at 25 °C. Following the reaction, the solution was concentrated to approximately 3 mL using a vacuum rotary evaporator. The target conjugates were isolated and purified by preparative thin-layer chromatography (prep-TLC) on silica gel plates. The plates were developed using a solvent system of dichloromethane/methanol (30:1, v/v) to successfully separate the Rapa-PEG conjugate from unreacted mPEG and coupling byproducts. The silica band containing the desired pure product was carefully scraped off and thoroughly extracted with ethyl acetate. The resulting suspension was filtered to remove the silica gel, and the filtrate was concentrated via a rotary evaporator before being completely dried in vacuo using a high-vacuum pump.

To monitor the purification and verify the product, Thin-Layer Chromatography (TLC) was performed. The purified product was spotted onto a silica TLC plate and developed in a chamber containing dichloromethane/methanol (20:3, v/v) as the mobile phase. Once the solvent front reached approximately 1 cm from the top, the plate was removed, air-dried, and subsequently stained in a chamber containing volatile iodine crystals to visualize the spots (Figure S4). The chemical structure and high purity of the resulting Rapa-PEG conjugates were comprehensively analyzed and confirmed by ^1^H Nuclear Magnetic Resonance (^1^H-NMR) spectroscopy (Bruker AVANCE III 400 MHz NMR spectrometer, utilizing CDCl_3_ as the solvent; **Figure S3**).

^1^H NMR (400 MHz, CDCl_3_) *δ* 6.45-5.1 (m, 10H, Rapamycin olefinic protons), 4.75-4.65 (m, 1H, Rapamycin C_40_-H esterified), 4.26 (t, *J* = 4.0 Hz, 2H, -CO-O-C**H_2_**-CH_2_-O-PEG), 3.61 (m, PEG backbone), 2.20-0.70 (m, Rapamycin aliphatic protons).

2. **Ultrasound Actuation Setup and Parameters**

A customized probe-type sonicator was utilized for all acoustic actuations. During the sonication process, the probe was vertically submerged approximately 1 cm below the air-liquid interface of the bacterial cultures, strictly avoiding contact with the vessel walls or bottom to prevent acoustic reflection and standing wave formation. During initial parameter optimization, acoustic frequencies of 5, 10, and 20 kHz were evaluated at varying amplitudes. A frequency of 20 kHz was selected for all subsequent in vivo gating and co-culture experiments due to its optimal balance: maximizing sonoporation efficiency for Rapa-PEG cellular entry while minimizing irreversible mechanical cell damage.

To prevent localized overheating and secondary thermal damage to the bacterial cells, a pulsed sonication paradigm was strictly adopted. Ultrasound was applied in intermittent duty cycles consisting of 1 minute of active sonication followed by a 3-minute recovery phase. Crucially, during this 3-minute resting period, the sample vessels were immediately returned to the orbital incubator and shaken continuously at 220 rpm. This intermediate shaking step was imperative for three reasons: (i) to rapidly re-homogenize the bacterial suspension, (ii) to effectively dissipate any localized heat generated near the probe tip, and (iii) to mechanically facilitate the regeneration and dissolution of gas microbubbles (cavitation nuclei) in the culture medium, which are rapidly depleted during active acoustic cavitation. The total active sonication duration varied from 0 to 10 minutes depending on the specific experimental design.

For the microbial consortium experiments involving two distinct strains (the "lysis" and "survivor" sub-populations), ensuring a uniform distribution of acoustic energy was exceptionally critical to prevent biased sonoporation or uneven growth stunting. Consequently, co-cultures were strictly handled in standardized vessels to maintain consistent acoustic field geometry. Throughout all acoustic treatments—whether for monocultures or co-cultures—the sample vessels were partially submerged in a temperature-controlled circulating water bath meticulously maintained at 35–37 °C. This meticulous thermal management ensured that the observed population dynamics and gene gating were solely attributed to the targeted chemo-acoustic triggering, completely decoupling any potential artifacts induced by cold shock or heat stress.

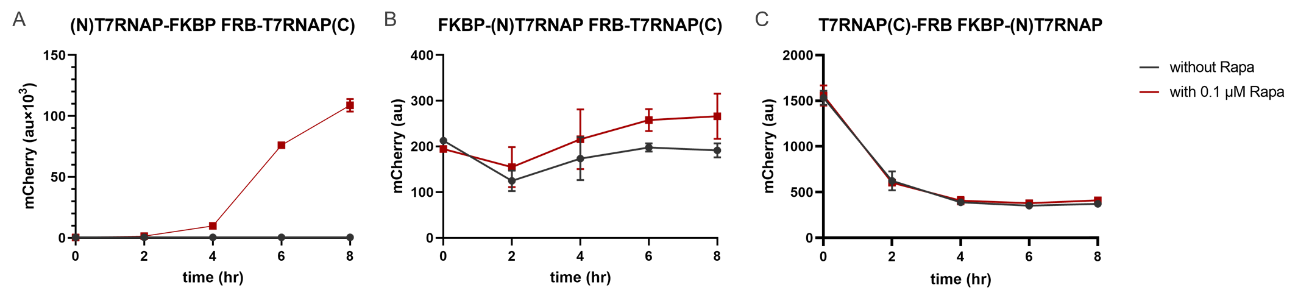

Figure S1: fusion orientations and linker compositions on the rapamycin biosensor performance with (A) N-terminal of T7RNAP fused with FKBP and C-terminal of T7RNAP fused with FRB; (B) FKBP fused with N-terminal of T7RNAP and FRB fused with C-terminal of T7RNAP; (C) C-terminal of T7RNAP fused with FRB and FKBP fused with N-terminal of T7RNAP.

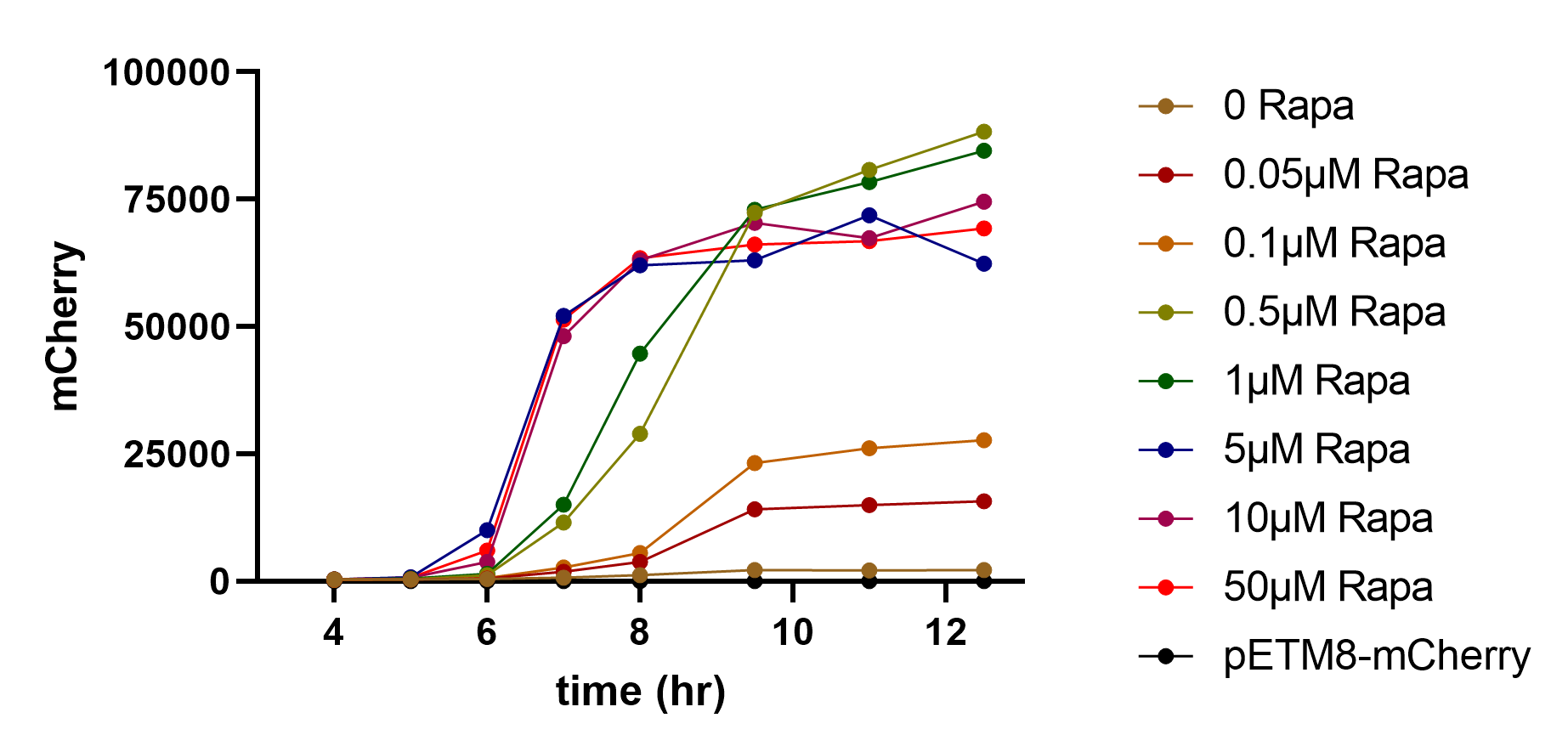

Figure S2: mCherry signal increasement over time induced by different rapamycin concentrations. Concentration above 10 $\mu$M did not further increase mCherry expression.

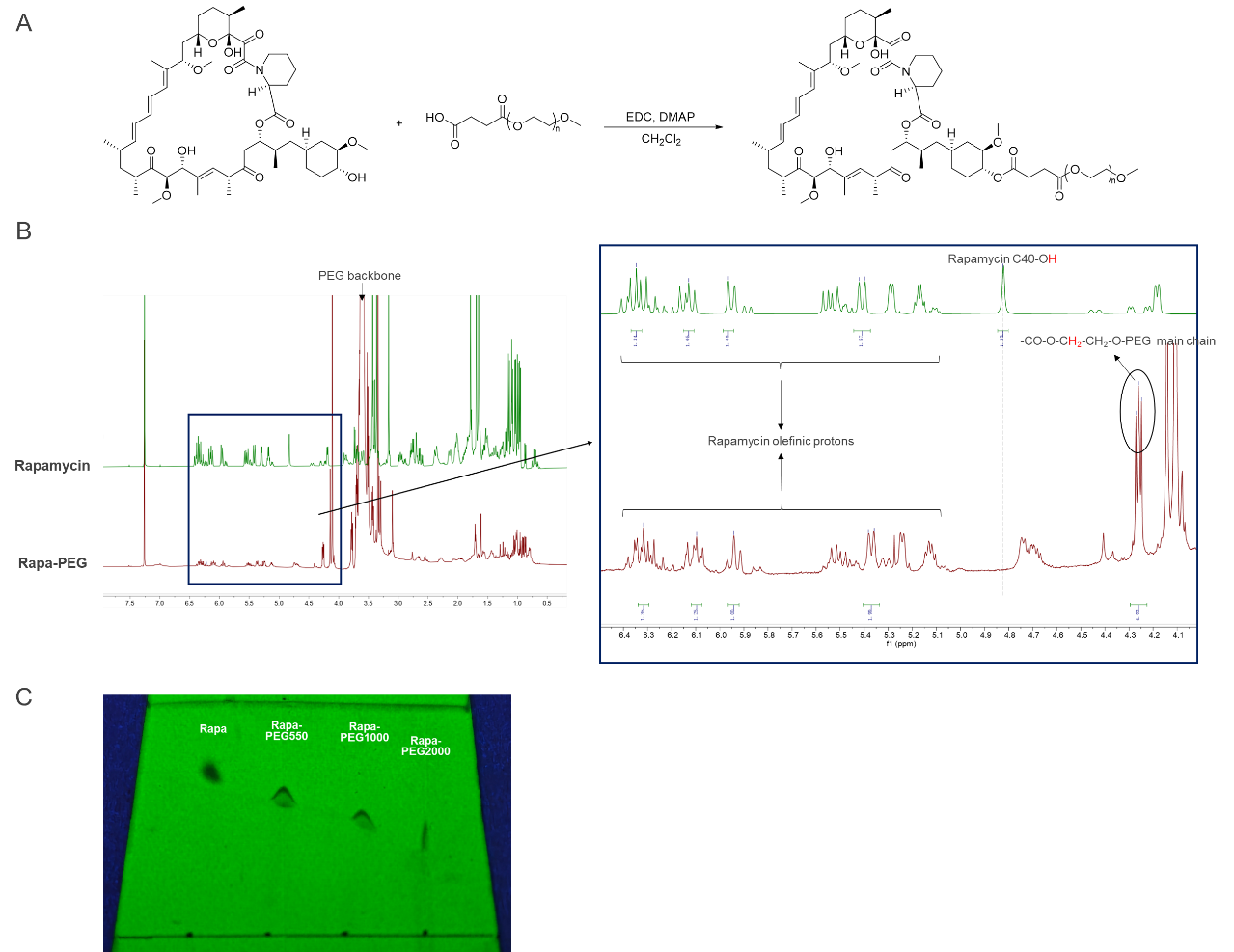

Figure S3: Synthesis and structural characterization of Rapa-PEG conjugates. (A) Synthetic scheme for the conjugation of Rapamycin to mPEG-COOH via an esterification reaction catalyzed by EDC and DMAP. (B) Stacked ¹H NMR spectra of free Rapamycin (green) and the synthesized Rapa-PEG conjugate (red) in CDCl₃. The inset shows a magnified view of the 4.15–4.35 ppm region. While free Rapamycin exhibits a weak native multiplet around 4.15–4.20 ppm, the Rapa-PEG conjugate displays a distinct, newly formed triplet at ~4.27 ppm. This triplet is unambiguously assigned to the methylene protons of the PEG linker immediately adjacent to the newly formed ester bond (-CO-O-**CH₂**-CH₂-O-). The minor signals observed at 4.12 ppm, along with 2.05 and 1.26 ppm, correspond to trace residual ethyl acetate used during the prep-TLC extraction process, which was entrapped by the viscous PEG chain despite high-vacuum drying. (C) Thin-layer chromatography (TLC) plate viewed under UV irradiation, visualizing free Rapamycin and Rapa-PEG conjugates with varying PEG chain lengths (Mw = 550, 1000, and 2000 Da).

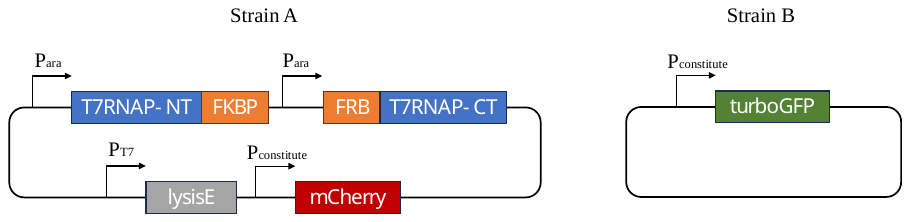

Figure S4: graphic illustration for strain A and B in co-culture system. Strain A carried the self-lysing motif under T7 promoter and mCherry under constitute promoter. Strain B carried turboGFP under the same constitute promoter.

Table S1: gene sequences of proteins used in this manuscript.

| **Gene** | **Sequence** |
| --- | --- |
| T7RNAP split at different positions | Atgaacacgattaacatcgctaagaacgacttctctgacatcgaactggctgctatcccgttcaacactctggctgaccattacggtgagcgtttagctcgcgaacagttggcccttgagcatgagtcttacgagatgggtgaagcacgcttccgcaagatgtttgagcgtcaacttaaagctggtgaggttgcggataacgctgccgccaagcctctcatcactaccctactccctaagatgattgcacgcatcaacgactggtttgaggaagtgaaagctaagcgcggcaagcgcccgacagccttccagttcctgcaagaaatcaagccggaagccgtagcgtacatcaccattaagaccactctggcttgcctaaccagtgctgacaatacaaccgttcaggctgtagcaagcgcaatcggtcgggccattgaggacgaggctcgcttcggtcgtatccgtgaccttgaagctaagcacttcaagaaaaacgttgaggaacaactcaacaagcgcgtagggcacgtctacaag (179/180)  aaagcatttatgcaagttgtcgaggctgacatgctctctaagggtctactcggtggcgaggcgtggtcttcgtggcataaggaagactctattcatgtaggagtacgctgcatcgagatgctcattgagtcaaccggaatggttagcttacaccgccaaaatgctggcgtagtaggtcaagactctgagactatcgaactcgcacctgaatacgctgaggctatcgcaacccgtgcaggtgcgctggctggcatctctccgatgttccaaccttgcgtagttcctcctaagccgtggactggcattactggtggtggctattgggctaacggtcgtcgtcctctggcgctggtgcgtactcacagtaag (302/303)  aaagcactgatgcgctacgaagacgtttacatgcctgaggtgtacaaagcgattaacattgcgcaaaacaccgcatggaaaatcaacaagaaagtcctagcggtcgccaacgtaatcaccaagtggaagcattgtccggtcgaggacatccctgcgattgagcgtgaagaactcccgatgaaaccggaagacatcgacatgaatcctgaggctctcaccgcgtggaaacgtgctgccgctgctgtgtaccgcaaggacaaggctcgcaagtctcgccgtatcagccttgagttcatgcttgagcaagccaataagtttgctaaccataaggccatctggttcccttacaacatggactggcgcggtcgtgtttacgctgtgtcaatgttcaacccgcaaggtaacgatatgaccaaaggactgcttacgctggcgaaaggtaaaccaatcggtaaggaaggttactactggctgaaaatccacggtgcaaactgtgcgggtgtcgataaggttccgttccctgagcgcatcaagttcattgaggaaaaccacgagaacatcatggcttgcgctaagtctccactggagaacacttggtgggctgagcaagattctccgttctgcttccttgcgttctgctttgagtacgctggggtacagcaccacggcctgagctataactgctcccttccgctggcgtttgacgggtcttgctctggcatccagcacttctccgcgatgctccgagatgaggtaggtggtcgcgcggttaacttgcttcct (563/564)  agtgaaaccgttcaggacatctacgggattgttgctaagaaagtcaacgagattctacaagcagacgcaatcaatgggaccgataacgaagtagttaccgtgaccgatgagaacactggtgaaatctctgagaaagtcaagctgggcactaaggcactggctggtcaatggctggcttacggtgttactcgcagtgtgactaagcgttcagtcatgacgctggcttacgggtccaaagagttcggcttccgtcaacaagtgctggaagataccattcagccagctattgattccggcaagggtctgatgttcactcagccgaatcaggctgctggatacatggctaagctgatttgggaatctgtgagcgtgacggtggtagctgcggttgaagcaatgaactggcttaagtctgctgctaagctgctggctgctgaggtcaaagataagaagactggagagattcttcgcaagcgttgcgctgtgcattgggtaactcctgatggtttccctgtgtggcaggaatacaagaagcctattcagacgcgcttgaacctgatgttcctcggtcagttccgcttacagcctaccattaacaccaacaaagatagcgagattgatgcacacaaacaggagtctggtatcgctcctaactttgtacacagccaagacggtagccaccttcgtaagactgtagtgtgggcacacgagaagtacggaatcgaatcttttgcactgattcacgactccttcggtaccattccggctgacgctgcgaacctgttcaaagcagtgcgcgaaactatggttgacacatatgagtcttgtgatgtactggctgatttctacgaccagttcgctgaccagttgcacgagtctcaattggacaaaatgccagcacttccggctaaaggtaacttgaacctccgtgacatcttagagtcggacttcgcgttcgcgtaa |
| FKBP | ATGGGCGTACAGGTGGAGACTATTTCCCCAGGCGATGGAAGAACATTTCCTAAACGTGGACAAACTTGTGTCGTCCATTACACTGGTATGCTTGAAGACGGTAAGAAGTTCGACAGTTCTCGCGACCGCAATAAGCCGTTTAAGTTTATGCTGGGGAAGCAAGAAGTTATACGCGGATGGGAAGAAGGTGTCGCTCAAATGAGTGTGGGGCAACGGGCGAAGCTGACCATTAGCCCAGACTACGCTTATGGTGCTACTGGGCACCCGGGCATAATTCCGCCTCATGCCACTTTGGTGTTCGACGTAGAACTTTTAAAGCTGGAGTAA |
| FRB | ATGGAGATGTGGCATGAAGGGTTGGAGGAGGCTTCACGGCTTTACTTCGGCGAGCGGAACGTCAAGGGAATGTTCGAAGTGTTGGAACCTCTTCATGCGATGATGGAAAGAGGCCCGCAGACACTGAAGGAGACCTCTTTTAACCAGGCTTATGGCCGGGATCTTATGGAAGCGCAGGAGTGGTGTCGCAAATATATGAAGAGTGGAAATGTCAAGGACCTGACCCAGGCTTGGGACCTTTACTACCACGTCTTTCGCCGGATTTCAAAGCAGTTACCCCAGCTTACCTCCTTAGAATTACAGTAA |
| lysis E codon optimized with LVA tag (in red) | ATGGTGCGCTGGACGCTGTGGGACACTTTGGCATTCCTTTTATTACTGAGTCTTTTGCTTCCCAGTCTGCTTATTATGTTCATTCCAAGTACGTTCAAGCGGCCAGTTTCCTCGTGGAAGGCGTTAAACCTGCGTAAGACCCTGTTAATGGCGTCTTCAGTGAGACTTAAACCCTTAAACTGTAGCAGACTGCCCTGTGTTTACGCGCAGGAAACGCTTACGTTCTTATTGACGCAAAAAAAAACGTGCGTGAAGAACTATGTTCAAAAAGAGGCCGCAAACGATGAAAATTATGCCCTGGTAGCGTAA |
| mCherry | atggtttcaaaaggcgaagaagacaacatggcgattatcaaggaatttatgcgtttcaaggtccacatggaaggcagcgtcaatggtcacgaatttgaaattgaaggcgaaggtgaaggccgtccgtatgaaggcacccagacggcaaaactgaaggtcaccaaaggcggtccgctgccgtttgcttgggatattctgtcaccgcaattcatgtatggttcgaaagcgtacgttaagcatccggccgatatcccggactatctgaaactgtcctttccggaaggcttcaaatgggaacgtgttatgaacttcgaagatggcggtgtggttaccgtcacgcaggatagctctctgcaagacggtgaatttatttataaagtgaagctgcgcggcaccaatttcccgagcgatggtccggttatgcagaaaaagacgatgggctgggaagcgagttccgaacgtatgtacccggaagacggtgccctgaaaggcgaaatcaagcagcgcctgaaactgaaggatggcggtcactatgacgcagaagtgaaaaccacgtacaaggctaaaaagccggtccaactgccgggtgcatacaacgtgaacatcaagctggatatcaccagccataacgaagactatacgatcgttgaacagtacgaacgtgcagaaggccgccactctaccggcggtatggatgaactgtacaaataa |
| turboGFP | ATGAGAGGATCGGGATCCGAGAGCGACGAGAGCGGCCTGCCCGCCATGGAGATCGAGTGCCGCATCACCGGCACCCTGAACGGCGTGGAGTTCGAGCTGGTGGGCGGCGGAGAGGGCACCCCCGAGCAGGGCCGCATGACCAACAAGATGAAGAGCACCAAAGGCGCCCTGACCTTCAGCCCCTACCTGCTGAGCCACGTGATGGGCTACGGCTTCTACCACTTCGGCACCTACCCCAGCGGCTACGAGAACCCCTTCCTGCACGCCATCAACAACGGCGGCTACACCAACACCCGCATCGAGAAGTACGAGGACGGCGGCGTGCTGCACGTGAGCTTCAGCTACCGCTACGAGGCCGGCCGCGTGATCGGCGACTTCAAGGTGATGGGCACCGGCTTCCCCGAGGACAGCGTGATCTTCACCGACAAGATCATCCGCAGCAACGCCACCGTGGAGCACCTGCACCCCATGGGCGATAACGATCTGGATGGCAGCTTCACCCGCACCTTCAGCCTGCGCGACGGCGGCTACTACAGCTCCGTGGTGGACAGCCACATGCACTTCAAGAGCGCCATCCACCCCAGCATCCTGCAGAACGGGGGCCCCATGTTCGCCTTCCGCCGCGTGGAGGAGGATCACAGCAACACCGAGCTGGGCATCGTGGAGTACCAGCACGCCTTCAAGACCCCGGATGCAGATGCCGGTGAAGAAGCCGCAAACGATGAAAATTATGCCCTGGTAGCGTGA |
| gapA constitute promoter | gtcgcaatgattgacacgattccgcttgacgctgcgtaaggtttttgtaattttacaggcaaccttttattcactaacaaatagctggtggaatat |

Table S2: primers used in this manuscript.

| **Primer** | **Sequence** |
| --- | --- |
| T7563-FKBP-T7fwd | tttaagaaggagatatacatatgaacacgattaacatcgctaagaa |
| T7563-FKBP-T7rvs | CGGAACCTCCaggaagcaagttaaccgcgc |
| T7563-FKBP-FKBPfwd | cttgcttcctGGAGGTTCCGGCGGGATG |
| T7563-FKBP-FKBPrvs | taattaagctgcgactagttttaCTCCAGCTTTAAAAGTTCTACGTCGA |
| T7563-FRB-FRBfwd | gttaacttgcttcctGGAGGTTCCGGCGGGGAGATGTG |
| T7563-FRB-FRBrvs | TAGGTtaattaagctgcgactagttttaCTGTAATTCTAAGGAGGTAAGCTGGGG |
| FRB-T7563-FRBfwd | tttaagaaggagatatacatatgGAGATGTGGCATGAAGGGTTGG |
| FRB-T7563-T7rvs | taattaagctgcgactagttttaaggaagcaagttaaccgcgc |
| FKBP-T7563-FKBPfwd | tttaagaaggagatatacatatgGGCGTACAGGTGGAGACT |
| FKBP-T7563-FKBPrvs | tcgtgttcatCCCGCCGGAACCTCCCTC |
| FRB-T7564-FRBrvs | cggtttcactCCCGCCGGAACCTCCCTG |
| FRB-T7564-T7fwd | TTCCGGCGGGagtgaaaccgttcaggacatct |
| FRB-T7564-T7rvs | aattaagctgcgactagttattacgcgaacgcgaagtccgact |
| FKBP-T7564-FKBPrvs | cggtttcactCCCGCCGGAACCTCCCTC |
| T7564-FKBP-T7fwd | tttaagaaggagatatacatatgagtgaaaccgttcaggacatct |
| T7564-FKBP-T7rvs | CGGAACCTCCcgcgaacgcgaagtccga |
| T7564-FKBP-FKBPfwd | cgcgttcgcgGGAGGTTCCGGCGGGATG |
| T7564-FRB-FRBfwd | cgcgttcgcgGGAGGTTCCGGCGGGGAGATGTG |
| 69-FKBP-T7rvs | CGGAACCTCCggcagcgttatccgcaa |
| 69-FKBP-FKBPfwd | taacgctgccGGAGGTTCCGGCGGGAT |
| FRB-70-FRBrvs | agaggcttggcCCCGCCGGAACCTCCCTGTAATT |
| FRB-70-T7fwd | TTCCGGCGGGgccaagcctctcatcactaccctac |
| 179-FKBP-T7rvs | CGGAACCTCCcttgtagacgtgccctacgc |
| 179-FKBP-FKBPfwd | cgtctacaagGGAGGTTCCGGCGGGAT |
| FRB-180-FRBrvs | taaatgctttCCCGCCGGAACCTCCCTGTAATT |
| FRB-180-T7fwd | TTCCGGCGGGaaagcatttatgcaagttgtcgaggc |
| FRB-303-FRBrvs | tcagtgctttCCCGCCGGAACCTCCCTGTAATT |
| FRB-303-T7fwd | TTCCGGCGGGaaagcactgatgcgctacgaagac |
| 302-FKBP-T7rvs | CGGAACCTCCcttactgtgagtacgcaccagc |
| 302-FKBP-FKBPfwd | tcacagtaagGGAGGTTCCGGCGGGAT |
| 600-FKBP-T7rvs | CGGAACCTCCctcatcggtcacggtaactactt |
| 600-FKBP-FKBPfwd | gaccgatgagGGAGGTTCCGGCGGGAT |
| FRB-601-FRBrvs | caccagtgttCCCGCCGGAACCTCCCTGTAATT |
| lysisE-fwd | aactttaagaaggagatatacatATGGTGCGCTGGACGCTGtgg |
| lysisE-rvs | aatATCGatgtcgatcctagCGCCAAGCTTcaaaaaacccctcaagaccc |
